# SARS-Cov-2-, HIV-1-, Ebola-neutralizing and anti-PD1 clones are predisposed

**DOI:** 10.1101/2020.08.13.249086

**Authors:** Yanfang Zhang, Qingxian Xu, Huikun Zeng, Minhui Wang, Yanxia Zhang, Chunhong Lan, Xiujia Yang, Yan Zhu, Yuan Chen, Qilong Wang, Haipei Tang, Yan Zhang, Jiaqi Wu, Chengrui Wang, Wenxi Xie, Cuiyu Ma, Junjie Guan, Shixin Guo, Sen Chen, Changqing Chang, Wei Yang, Lai Wei, Jian Ren, Xueqing Yu, Zhenhai Zhang

## Abstract

Antibody repertoire refers to the totality of the superbly diversified antibodies within an individual to cope with the vast array of possible pathogens. Despite this extreme diversity, antibodies of the same clonotype, namely public clones, have been discovered among individuals. Although some public clones could be explained by antibody convergence, public clones in naïve repertoire or virus-neutralizing clones from not infected people were also discovered. All these findings indicated that public clones might not occur by random and they might exert essential functions. However, the frequencies and functions of public clones in a population have never been studied. Here, we integrated 2,449 Rep-seq datasets from 767 donors and discovered 5.07 million public clones – ~10% of the repertoire are public in population. We found 38 therapeutic clones out of 3,390 annotated public clones including anti-PD1 clones in healthy people. Moreover, we also revealed clones neutralizing SARS-CoV-2, Ebola, and HIV-1 viruses in healthy individuals. Our result demonstrated that these clones are predisposed in the human antibody repertoire and may exert critical functions during particular immunological stimuli and consequently benefit the donors. We also implemented RAPID – a **R**ep-seq **A**nalysis **P**latform with **I**ntegrated **D**atabases, which may serve as a useful tool for others in the field.

## Background

Antibody is a critical immunoglobulin complex consisting of two identical heavy and two identical light chains. Each chain is encoded by selectively recombining one of the various germline gene fragments, namely variable (V), diversity (D, for heavy chain only), and joint (J) genes. The sequence between V gene end and J gene start is called complementarity determining region 3 (CDR3) which is extremely diverse because of the random nucleotide insertion and deletion in the junctions and by large defines the binding specificity of an antibody^1^. This binding specificity makes antibodies favorable for therapeutic purposes.

With tremendous efforts and various techniques, many monoclonal antibodies (mAbs) targeting distinct viruses and proteins were discovered in the past decades^2,3^. However, the primary barrier to studying antibodies is their immense diversity. The total number of antibodies in an adult, termed antibody repertoire, is estimated to be around 10^12^ – a number far out of reach for these traditional methods^4^.

Fortunately, antibody repertoire sequencing (Rep-seq) was invented to acquire millions of antibody variable regions in DNA or RNA form in a single experiment, a great advance thanks to the advent of high-throughput sequencing (HTS) technology. With the aid of this technique, our understanding of the humoral immunity was markedly advanced and many valuable mAbs were identified. For instance, we and others have used Rep-seq method to discover HIV-1 broad neutralizing antibodies^5–7^. It also helped researchers in identifying neutralizing antibodies in the recent SARS-CoV-2 outbreak^8–10^. Thus far, Rep-seq has been proved to be productive in studying cancer immunology^11^, virus infection^12,13^, vaccination^14,15^, etc.

Besides the achievement aforementioned, this data-rich method also led to the finding of public clone – antibodies in different individuals but share the same or similar CDR3 which implies the same binding specificity. The fraction of public clones between two individuals is estimated to be ~0.95% to 6%^16,17^ in circulating repertoire, or linearly correlate with the product of total clones of the sample pair^18^. They were found in individuals infected with the same virus and thus implicated the antibody convergence, a phenomenon in which antibodies are assimilated to each other^19^. Later, they were also present in B1 and marginal zone B cells in the naïve state20. For instance, Soto *et al*. revealed public clones in cord blood^16^. Intriguingly, Jardine *et al.* found VRC01-class HIV-1 neutralizing antibody clones in naïve B cells from healthy individuals^21^. Kreer *et al.* discovered SARS-CoV-2-neutralizing clones in uninfected healthy people^8^. All these studies were conducted on a limited number of samples, the answers to some of the key questions about public clone remained unsolved. What proportion of an antibody repertoire is public at a population level? What are the other public clones existing in the human repertoire? Have they undergone maturation process? What are their functions? Do they influence our health during disease onset or virus infection?

Bearing these questions in mind, we collected 88,059 known antigen-binding or disease-associated antibodies published before, 521 therapeutic antibodies recorded by the World Health Organization (WHO)^22^, and 2,449 high-quality Rep-seq datasets (767 donors, 306 million clones, and 7.12 billion raw reads) published by others as well as generated in our lab. Integrative and systematic analysis revealed that there are around ~ 10% or more public clones for each individual in a population level. Three thousand three hundred and ninety of these public clones can be annotated indicating they are functionally important for humoral immune response. More importantly, we found public clones that binding to PD1, neutralizing SARS-CoV-2, Ebola, and HIV-1 viruses in healthy individuals. These results demonstrated that public clones in the population are predisposed in the repertoires of particular individuals who may later benefit from their existence upon virus infections and disease onset. All datasets in this study were integrated and implemented in RAPID – <u>**R**</u>ep-seq <u>**A**</u>nalysis <u>**P**</u>latform with <u>**I**</u>ntegrated <u>**D**</u>atabase, a knowledge-rich platform for others to analyze and annotate their own repertoire data.

## Result

### RAPID: a powerful platform for Rep-seq data analysis

Currently, a substantial number of tools or web servers have been proposed to address the issues of Rep-seq data analysis or characterization for repertoires^23–40^. However, these platforms focus on analyzing Rep-seq dataset individually and ignoring exploration of discriminating repertoire features within or among groups. Apart from that, antibody databases are also specialized for antigen annotation, such as bNAber which just documents HIV broadly neutralizing antibodies^41^. Thus, our platform named RAPID which compensates for shortages above was built. As shown in Fig. 1a, the data repositories comprised three different data modules, namely Rep-seq data collection, therapeutic antibody collection, and known antibody collection. The Rep-seq data integrated 2,449 high-quality datasets (see Yang *et al.* for method18) from 767 donors either downloaded from published data repository or generated in our lab. These datasets contain samples from different genders, various tissues, immune status, and age spans, and were generated via different amplification strategies. Thus, it provided a rich source of reference for analyzing and comparing antibody repertoires. There are 7.12 billion reads and 306 million clones yielded from a systematic analysis pipeline using exactly the same criteria, thus making them comparable to each other^18^. The therapeutic monoclonal antibodies (mAbs) were downloaded from the Thera-SAbDab database which contains 521 therapeutic mAbs of different types at various stages. The 88,059 known antibodies were downloaded from multiple data repositories and carefully annotated via natural language processing method as well as manual check (Supp. Fig. 1 and Materials and Methods). These annotations included antibody sequences of different chain types as well as the antibodies that binding to and neutralizing virus, associating to particular diseases, etc. The raw sequences, CDR3s, descriptions, and sample metadata information were systematically extracted and stored in a rational database as well as FASTA format files when necessary. A user-friendly interface for searching particular terms, antibodies, and CDR3s (Supp. Fig 2) was also provided (https://rapid.zzhlab.org/).

**Figure 1.**
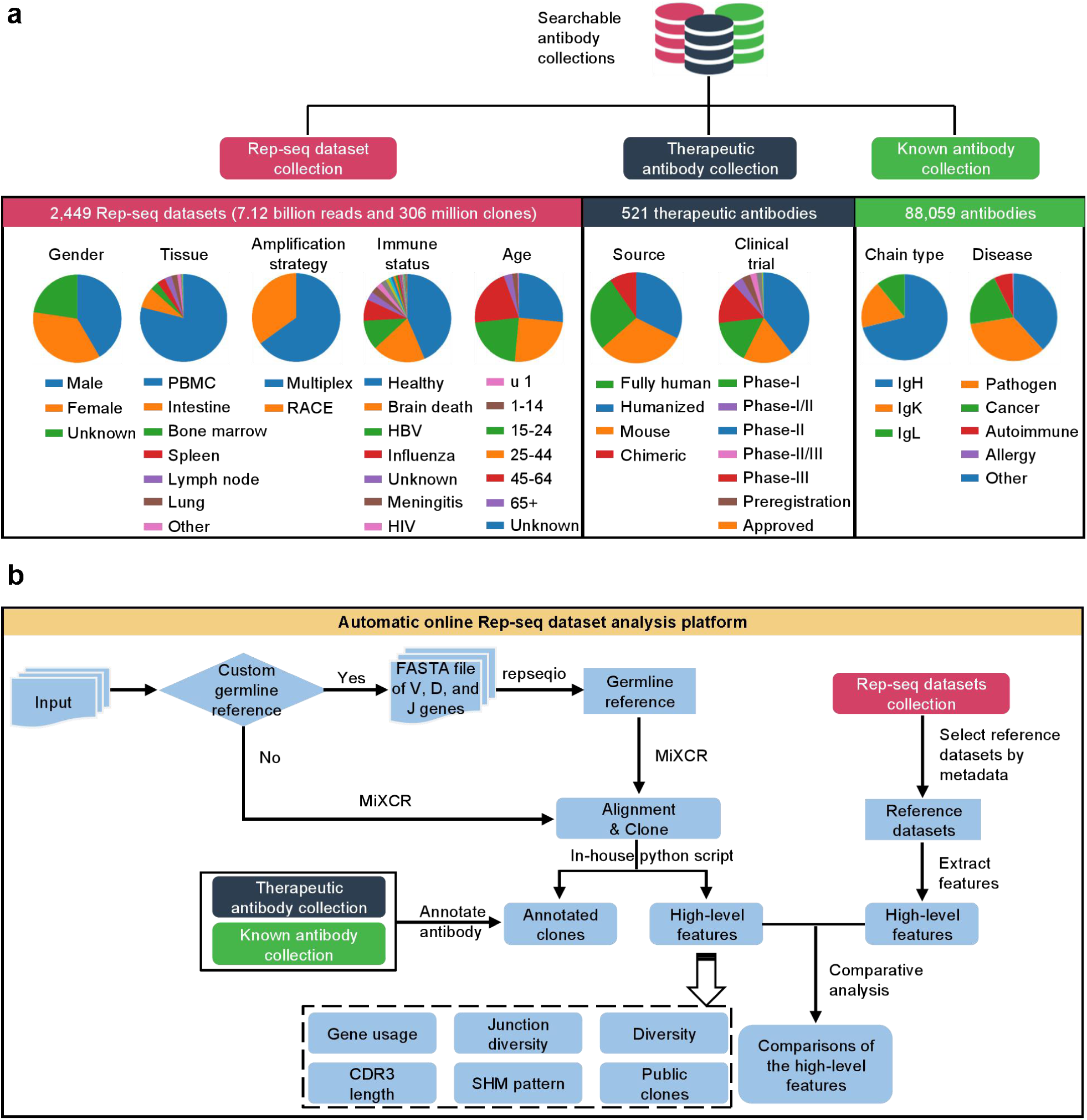
Data composition and automatic online Rep-seq dataset analysis pipeline of the RAPID. **(a)** Composition and detailed information of the three types of antibody datasets: Rep-seq datasets, therapeutic antibodies and known antibodies. The pie charts in the lower panel showed detailed composition of each type of antibodies. **(b)** The analysis platform for Rep-seq dataset.

**Figure 2.**
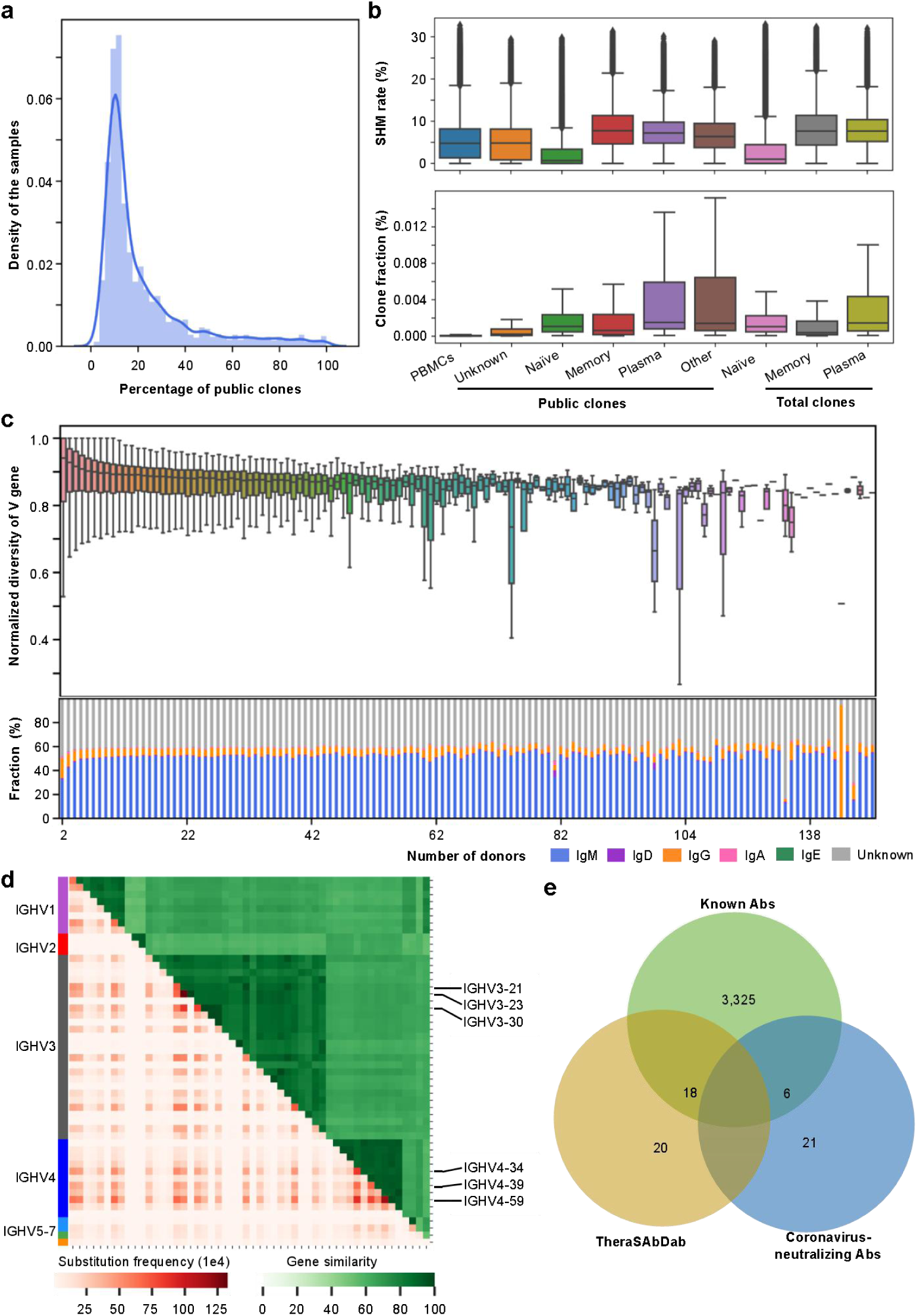
The characteristics of public clones. **(a)** The sample density distribution with regard to the fractions of public clones. X-axis represents the percent of public clones in a sample. Y-axis shows the sample density. **(b)** Comparisons of the distribution of somatic hypermutation rates (SHMs, upper panel) and clone fractions (lower panel) among different groups of clones. **(c)** Upper panel: The normalized Shannon index of V gene usage (Y-axis) within each CDR3 aa group sorted by the respective numbers of shared donors (X-axis). Lower panel: The stacked bar plot showed the composition of public clones for different antibody isotypes. **(d)** The substitution frequencies (lower left panel) and percent identities of V genes (upper right panel). **(e)** The overlap of 3,390 annotated public clones with known antibodies, thera-SAbDab, and Coronavirus-neutralizing antibodies.

The data analysis module allows users to streamline their own data through a versatile pipeline. Apart from the general low- and high-level analyses, the RAPID also provides some helpful features as described below (Fig. 1b). 1) Customizing germline reference. 2) Customizing reference datasets; the users can freely select one or more datasets in the platform as reference for cross-comparisons purpose. 3) Automatic antibody annotation; the CDR3s from the input dataset will be automatically compared to the CDR3s in the data repository on RAPID and annotated where applicable. 4) Downloadable figures and analysis result. Thus, any researcher can upload their datasets and cross compare them to 2,449 datasets with 306 million clones, all therapeutic antibodies, and 88,059 known antibodies and retrieve the relevant information. With the thorough antibody collections and a versatile analysis platform, we believe the RAPID will be helpful for the large cadre of scientists who demand analyzing antibody repertoire data as we demonstrated in this study.

### Public clones are prevalent in the population

With this unprecedented dataset, we started in-depth inspection of the public clones. Here, we defined public clones as antibodies with the same CDR3 amino acid sequence that present in more than one donor. Even with this stringent criterion, we discovered 5,077,372 public clones. Typically, the public clones represent ~1.23 to 100 percent of each individual repertoire with a peak value of 10.46 percent (Fig. 2a). Moreover, 65 public clones occur in more than 100 individuals with one clone shared by 196 individuals (25.55% of the total donors) (Supp. Fig. 3 a and b). Thus, population level study helped us find more public clones and highly frequent ones. As 96.86% of the public clones are from PBMCs (Supp. Fig. 3c), we compared their SHMs and clone fractions of the clones in different groups. The SHMs for naïve, memory, and plasma groups were comparable between public clones and total clones. The public clones of PBMCs and unknown samples displayed mediocre SHMs between naïve and non-naïve clones (Fig. 2b, upper panel). For clone fractions, the public clones from PBMCs and unknown samples were lower than the other counterparts (Fig. 2b, lower panel) indicating they are inactivated. On the other hand, about half of the public clones were from IgM isotype (Fig. 2c, lower panel). Therefore, it’s reasonable to speculate that majority of these public clones were acquired from naïve and lowly-mutated memory B cells.

**Figure 3.**
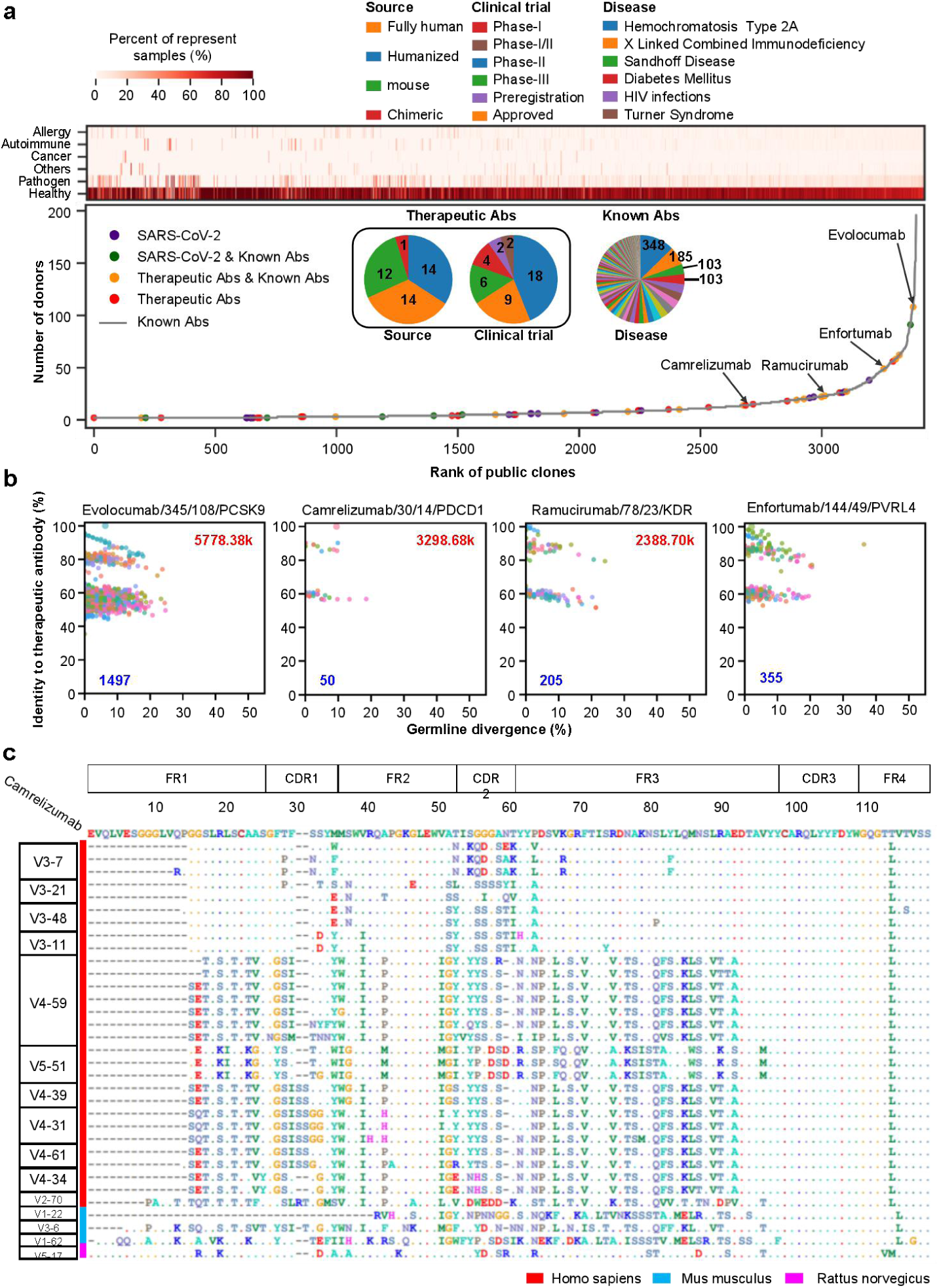
The characteristics of annotated public clones. **(a)** Overview of 3,390 annotated public clones. The upper heatmap stand for the composition of samples for these annotated public clones. Samples were divided into 6 groups including allergy, autoimmune, cancer, pathogen, healthy, and others. The bottom line chart means the number of donors. Public clones annotated by known antibody database were shown by gray line, while those annotated by other databases were shown by scatters filled in particular colors. The center top pie charts show distribution of source and clinical trial for therapeutic antibodies and associated disease for known antibodies. **(b)** Identity of variable region sequences from FR1 to FR3 of antibodies whose CDR3aa are same as Evolocumab, Camerelizumab, Ramucirumab, and Enfortumab. The X-axis means the germline divergence and different colors of scatters mean different V genes. Numbers filled in red stand for the death caused by diseases treated by such antibody and those filled in blue stand for the number of variable regions identified from Rep-seq datasets. Titles for sub-figures separated by forward slash include inn id of therapeutic antibody, the number of samples and donors with such CDR3aa, and target of therapeutic antibody. Dots for therapeutic antibodies are larger than that of clones identified from Rep-seq datasets. **(c)** Multiple sequences alignment of variable region for antibodies with the same CDR3aa as Camerelizumab from homo sapiens, mus musculus, and rattus norvegicus. The amino acid sequence of Camerelizumab is listed in the top and regions (FR1-FR4) for variable region are marked by boxes above. V gene used in each sequences are labeled in left boxes.

We also observed that different V and J gene combinations can yield the same CDR3s. As expected, the diversity of V gene usage for public clones increased when clones are shared by more donors (Supp. Fig. 4). However, when normalized to the maximum theoretical diversity with particular number of V genes (see Materials and Methods), this diversity slightly decreased with the increment of sharing donors indicating recombination preference of V genes (Fig. 2c, upper panel). Careful examinations of the V and J genes that formed the same CDR3s showed that same J gene was always preferred while V gene was more replaceable among individuals (Fig. 2d and Supp. Fig. 5). Nevertheless, the substitution rates of V genes were not completely influenced by their sequence similarity (Fig. 2d). This result suggested that J genes might affect the CDR3s more than V genes do.

**Figure 4.**
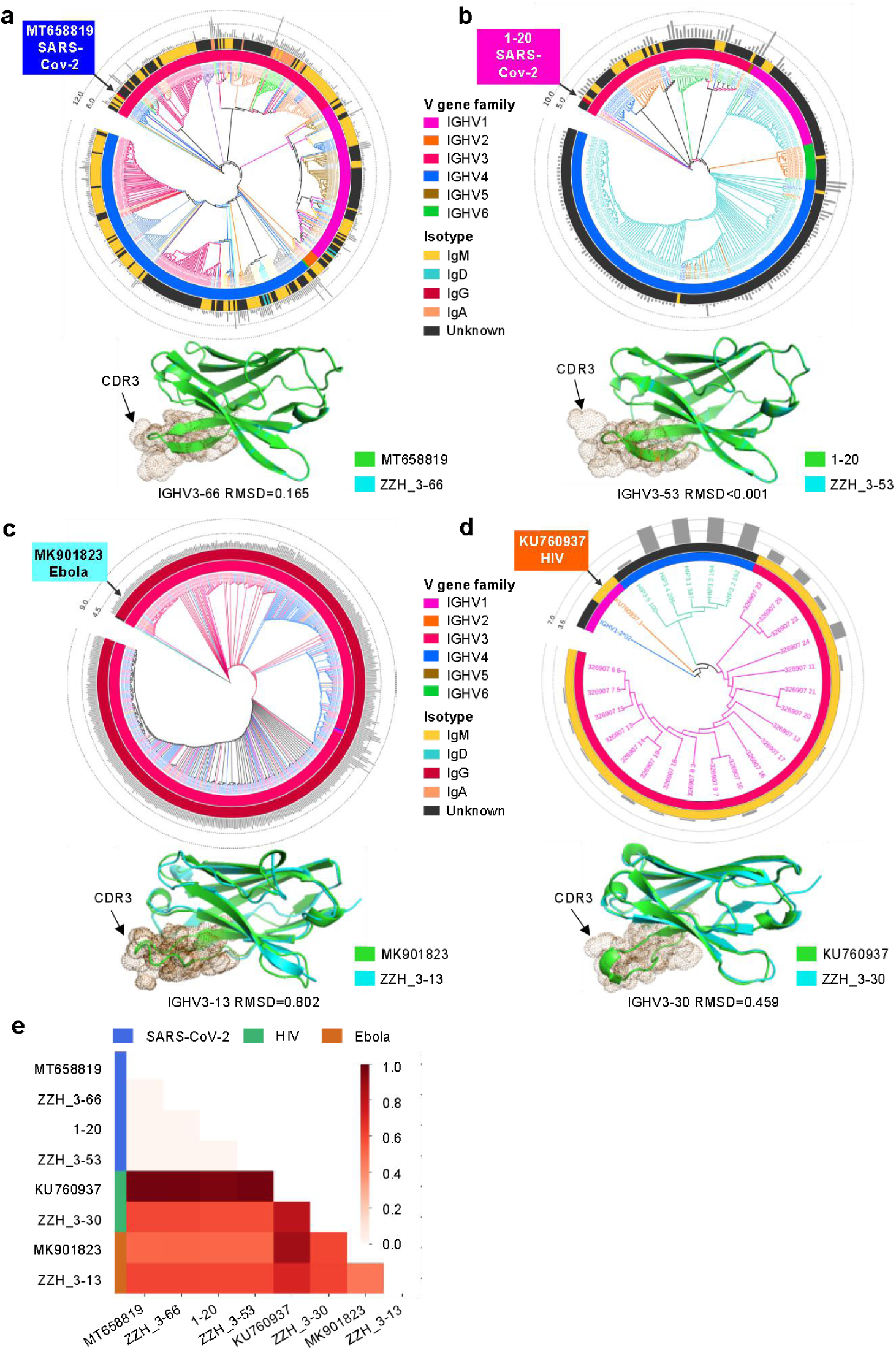
Maturation pathway and structure of antibodies potentially targeting SARS-CoV-2, Ebola, and HIV. Variable region sequences with the same CDR3aa as MT658819 **(a)**, ^1–20^ **(b)**, MK901823 **(c)**, and KU760937 **(d)** were extracted and compared with those validated neutralizing antibodies and their germline reference. The germline reference was chosen as root of phylogenetic tree and the validated antibodies were marked by arrow. The cluster map contains four layers including similarity of sequences (the sequences extracted from the same donor were marked with the same color), V gene family, isotype, and somatic hypermutation rate from inner to outer. Under each phylogenetic tree, similarity of structures for validated antibody and variable region identified from Rep-seq datasets were shown. **(e)** Pair-wise structure comparison of antibodies targeting SARS-CoV-2, HIV, and Ebola.

Taken advantage of the rich antibody information integrated in this study, we tried to annotate these public clones. Totally, 3,390 public clones have been annotated by three antibody databases including known antibody, Thera-SAbDab, and Coronavirus-neutralizing antibody incorporated with 459 mAbs from CoV-AbDab^42^, 28 mAbs from Kreer *et al.*^8^, and 19 mAbs from Liu *et al.*10 (Fig. 2e). We found that 3,349 out 3,390 clones shared the same CDR3s amino acid sequences with known antibodies targeting specific antigens or associating with diseases. Surprisingly, we found 38 and 27 CDR3s overlapping with Thera-SAbDab and Coronavirus-neutralizing antibodies. Among 27 Coronavirus-binding clones, 8 of them target SARS-CoV-2 (CARGDSSGYYYYFDYW binds both SARS-CoV-1 and SARS-CoV-2). As all these Rep-seq datasets were generated before the outbreak of the COVID-19, these SARS-CoV-2-neutralizing clones and therapeutic clones were deposited in the antibody repertoire. These provide evidence that those therapeutic and antigen-specific antibodies exist widely in populations and can be a powerful source for mAb discovery with clinical purposes.

### Therapeutic mAb clones are prevalent in healthy people’s repertoire

We went on to look into the details of the 3,390 annotated public clones. Among them, 3,354 (98.94%) were found in at least one healthy sample (Fig. 3a, upper panel). The therapeutic antibodies found in this data collection compromise 41 therapeutic mAbs with 38 unique CDR3s. Of these, six are under phase III clinical trial, two are under preregistration and nine are approved for clinical usage (Fig. 3a, lower panel). Interestingly, only 14 (34.15%) mAbs are fully human while the others include 14 (34.15%) humanized, 12 (29.27%) from mouse, and one chimeric. This indicated that therapeutic antibodies generated by mouse model can be generated by our own immune system. Thus, public clone might be a better source for mAb discovery for clinical usage.

Also, many of the therapeutic antibody clones found in healthy people which are used for treatment of diseases with top causes of death in the world (Table 1) prevailed in the population (Fig. 3, a and b). For instance, Evolocumab targeting PCSK9 is used to treat Coronary disorders, Stroke, and Hypercholesterolaemia. Stroke alone caused 5.78 million deaths worldwide in 201643 and this clone was found in 108 (14.1% of the total of 767) donors’ repertoire. The CDR3 of anti-PD1 (Camrelizumab), the treatment to various cancers, was found in 14 donors’ repertoire. Ramucirumab targeting KDR and Enfortumab targeting PVRL4 were also found in 23 and 49 donors, respectively. According to the percent identities to therapeutic antibodies, most of the antibodies from the same clonotype separated into at least two groups (Fig. 3b and Supp. Fig. 6). Detailed inspection revealed that multiple V genes involved in the recombination, again supported the diversity of V genes within clones (Fig. 2d). These antibodies might serve as therapeutic alternatives for the same disease.

**Table 1.**
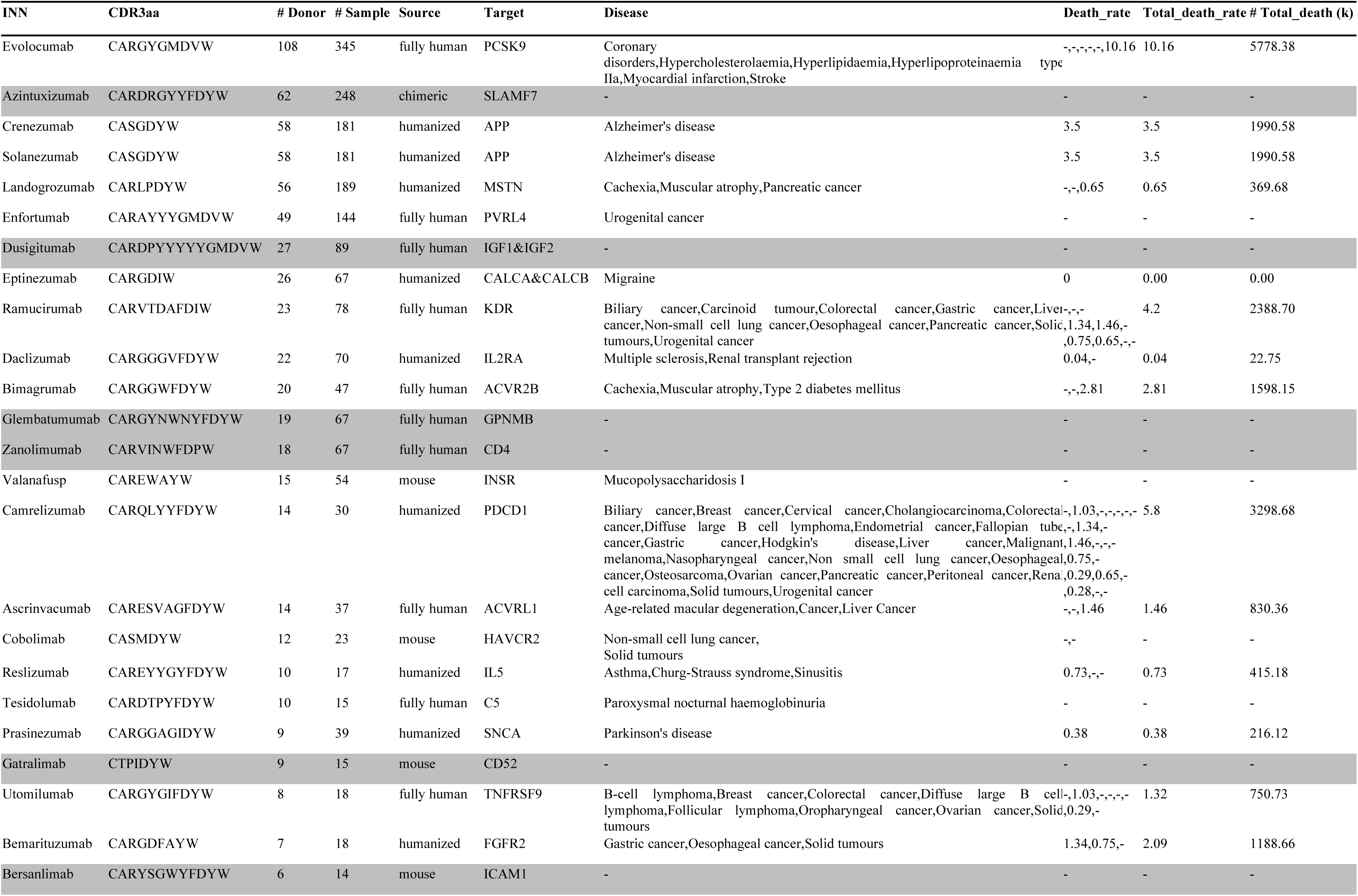
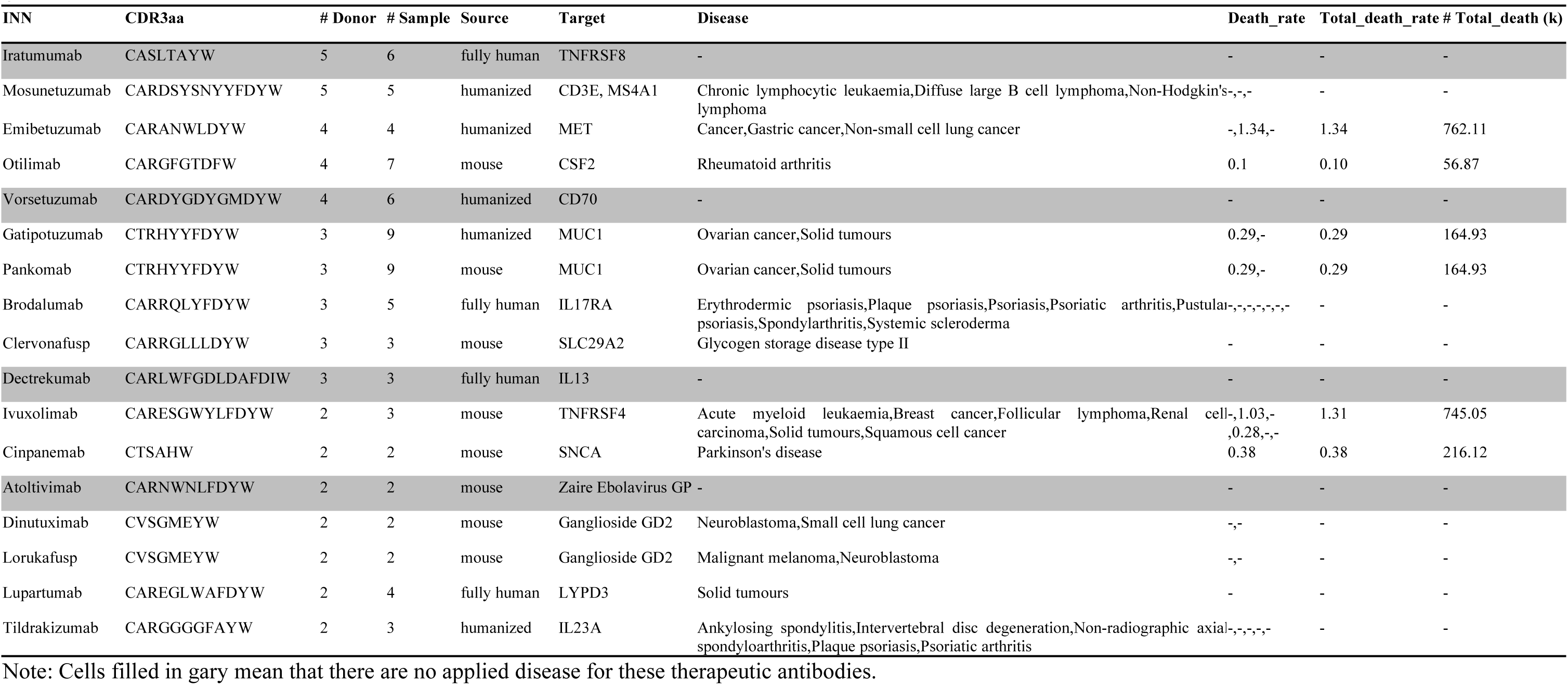
Detailed information of therapeutic antibodies whose CDR3aa were same as those identified 232 from Rep-seq datasets

Moreover, for therapeutic clones, there were always Rep-seq captured antibodies showing higher SHM rates (Fig. 3b), possibly indicated the engagement of maturation process under the selective pressure of their antigens.

Interestingly, we also found antibodies with the same CDR3s as anti-PD1 antibody in the repertoires of mouse and rat. The variations in the other regions of variable sequences (Fig. 3c) indicated that they were purposely selected and retained in the repertoire. Moreover, the differences in their structure might be the result of structural variations of PDCD1 proteins in different species (Supp. Fig. 7). Finding anti-PD1 antibody clones in healthy individuals and other species was exciting. On the contrary, no anti-PD1 clone was found in any of the 56 samples of various cancers – 2 breast cancer, 49 colorectal cancer, and 5 liver cancer samples. Why cancer patients do not possess anti-PD1 antibody clones is an intriguing question. Understanding the mechanism behind this fact may lead to a better understanding of cancer immunology and more effective immunological therapy. Besides, further studies illustrating the causes and consequences of the anti-PD1 clones in particular individuals would also benefit the field.

### SARS-CoV-2-, Ebola-, and HIV-1-neutralizing antibody clones are predisposed

On a par with the results in finding therapeutic clones, multiple clones neutralizing SARS-CoV-2, Ebola, and HIV-1 were also uncovered from virus-naïve individuals. For SARS-CoV-2-neutralizing clones MT658807, MT658819, and ^1–20^, there were 20, 506, and 222 heavy chain variable regions extracted from repertoires of 5, 17, and 6 healthy donors, respectively. Twenty-five variable regions from 2 healthy donors sharing the same CDR3 sequence with HIV-1-neutralizing class VRC01 were also extracted. In addition, 2,663 variable regions from 4 donors injected with influenza vaccine contained the CDR3 of Ebola-neutralizing antibody of MK90182344. Although neutralizing clones to HIV-1 and SARS-CoV-2 were reported to exist in the repertoire of the naïve B cell previously^8,21^, this is the first time to identify these neutralizing clones in multiple people. Thus, we concluded that they are predisposed in a population.

We then set off to explore the maturation pathways of these neutralizing clones by analyzing the phylogenetic trees of each clone built via DNAMLK (Fig. 4a-d). The Ebola-neutralizing clones exhibited high maturation rates with IgG. Interestingly, the maturation rates of three SARS-CoV-2 clones demonstrated various level of SHMs. While the overall SHM rate for MT658807 clone is lower than 2.5%, some of the antibodies in MT658819 and ^1–20^ clones displayed more than 5% mutations. Previous studies reported the general lower SHM for SARS-CoV-2-neutralizing antibodies but still some clones with more mutations were identified and verified^8–10^. Therefore, different clones might subject to different selective pressure and consequently manifest various SHM rates. Although the diversity of V gene usage in neutralizing clones by large defined the topology of the phylogenetic trees, oftentimes antibodies from different individuals aggregating in the same branch was observed. This indicated antibody convergence in the same maturation pathway.

Apart from the MK901823 clone, which was from a sample after influenza vaccine trivalent, inactivated seasonal influenza (TIV), all the donors of the HIV-1 and SARS-CoV-2 clones are virus-free healthy individuals. Furthermore, we found the same antibodies of MT658807 in donor 1776. This again confirmed the predisposition of this neutralizing clone. However, what triggers their maturation would be an important question to answer for future studies.

To further explore their possibility of virus binding, we compared the structures of these Rep-seq retrieved antibodies to their corresponding verified neutralizing antibodies. As shown in Fig. 4a-d, we found antibodies were very similar to MT658819^8^ (RMSD: 0.165) and 1-20^10^ (RMSD: < 0.001) that neutralizing SARS-CoV-2, MK901823^44^ (RMSD: 0.802) that neutralizing Ebola, and KU760937^21^ (RMSD: 0.459) that neutralizing HIV.

To validate this similarity, we performed pair-wise structure comparison among antibodies the neutralizing these three viruses. As shown in Fig. 4e, the RMSD scores of clone targeting the same antigen were much lower than those targeting different antigens. Thus, the high similarity of Rep-seq retrieved antibodies to neutralizing antibodies are reliable.

### SARS-CoV-1-neutralizing and therapeutic antibody clones exist in animals

Inspired by the existence of anti-PD1 clones in mouse and rat, we scrutinized the Rep-seq datasets with four different species, namely Macaca fascicularis, Macaca mulatta, Mus musculus, and Rattus norvegicus. We found 4 SARS-CoV-1-neutralizing and 18 therapeutic clones in at least one species. Taken together, we believe these clones are not randomly generated but purposely selected and disposed in vertebrates’ repertoire.

## Discussions

Public clones are a specific fraction of antibodies among individuals that we know little about. By integrating the largest antibody data to date, population level analyses discovered millions of public clones which represent ~10% or higher fraction of each individual’s repertoire. However, compared to the superb diversity of the antibody repertoire, the current dataset might still be smaller than demand. We believe that when more datasets will be integrated, more public clones would be revealed. This is understandable since although the somatic recombination may generate numerous antibodies, majority of them are eliminated during the negative selection process in the bone marrow. Consequently, the once private repertoire might be public^45^.

How often can we find these public clones with critical functions in an individual? Are they predisposed in everyone’s repertoire? The current data seems to support that only some people possess them. However, we found that sequencing depth is critical for public clone identification as many more public clones were observed in datasets with very high depth. Currently, only a few hundred thousand to a few million reads were captured in general. Compared to the theoretical number of B cells in the sample and the depth needed to identify a clone confidently, much more sequencing reads are demanded. As most of the therapeutic mAbs target proteins of conserved genes such as PDCD1, another helpful practice in finding functional public clones might be comparing antibody repertoires between human and other vertebrates.

The finding of clones that can bind to PDCD1 or neutralize SARS-CoV-2, Ebola, and HIV-1 viruses demonstrated that public clones might be important for the donor’s health. Then discovering the functionalities of the vast majority of other public clones would be critical for a deep understanding of the humoral immune system. The major challenge in this regard is the lack of the light chain pair. The techniques of paired heavy and light chain sequencing invented in Georgiou lab^46^ and the single cell repertoire sequencing^47^ showed great potential in solving this problem.

We’ll update RAPID along with the accumulation of Rep-seq datasets generated by others and our lab. We believe more public clones will be identified and their functions will be illustrated along this path.

## Methods

### Rep-seq datasets enrollment

Method to enroll published and in-house Rep-seq datasets were described in Yang *et al*^18^, please refer to it for detailed information. The re-analysis pipeline of these Rep-seq datasets was also included in that paper.

### Resources of known antibody

Five open access antibody databases, named abYsis (http://abysis.org/)^48^, bNAber (http://bnaber.org/)^41^, EMBLIG (http://acrmwww.biochem.ucl.ac.uk/abs/abybank/emblig/), HIV Molecular Immunology Database (HIV-DB: https://www.hiv.lanl.gov/content/immunology/neutralizing_ab_resources.html)^49^ and IMGT/LIGM-DB (http://www.imgt.org/ligmdb/)^50^ were enrolled. In addition, another two nucleotide sequence databases, including European Nucleotide Archive (ENA) of EMBL-EBI (https://www.ebi.ac.uk/ena/data/view/Taxon:9606&result=coding_release)^51^ and National Center for Biotechnology Information (NCBI) Nucleotide database (https://www.ncbi.nlm.nih.gov/nucleotide/), were also incorporated. In a word, 7 databases were finally included. The search strategy and download date for them were listed in Supplementary Table 1.

### Construction of known antibody database

Although the species was restricted for sequences downloading, some sequences from other species, like Mus musculus, were also included. Thus, we firstly discarded non-human sequences according to descriptions. After that, sequences were aligned to V, D, and J germline reference (downloaded from IMGT: http://www.imgt.org/ and listed in Supplementary Table 2) by IgBLAST^24^ (version 1.8.0), as its great performance for error-corrected reads^23^. Based on results of IgBLAST, sequences which meet criteria were reserved, including in-frame, productive, with V, J, and CDR3, without either stop codon or out-of-frame in variable region, and without ambiguous base (N) in CDR3. Sequences with the same nucleotide sequences of variable region within the same database were de-duplicated. To remove non-antibodies from NCBI and ENA, we aligned these sequences to NCBI Nt database (downloaded from NCBI: ftp://ftp.ncbi.nlm.nih.gov/blast/db/FASTA/nt.gz) and discarded the sequences whose descriptions contain no keywords we defined (Listed in Supplementary Table 3). These keywords were selected from descriptions of the antibody sequences are stored in the database of abYsis, bNAber, HIV-DB, EMBLIG, and IMGT/LIGM-DB. Furthermore, antibodies from 7 databases were pooled together and de-duplicated according to the nucleotide sequence of variable region. In the end, disease information for antibodies from EMBLIG, ENA, IMGT/LIGM-DB, and NCBI was annotated by TaggerOne^52^ based on description, title, and abstract of sequences. The sequences from abYsis were annotated as “NA”, as no annotation information can be downloaded. The sequences from HIV-DB and bNAber were annotated as HIV infections.

### Implementation of RAPID

The web interface is implemented by Hyper Text Markup Language (HTML), Cascading Style Sheets (CSS), and JavaScript (JS). It is a single page application based on the JS framework React.js, while using the React component library Ant Design to unify the design style. The back end of the website uses Nginx as the HTTP and reverse proxy server, develops business logic based on Node.js, uses MySQL to manage data, and uses RabbitMQ to process the analysis task queues. Furthermore, the real-time notification of task progress depends on the WebSocket technology.

### Extraction of variable region identified from Rep-seq dataset

Firstly, if regions from FR1 to FR4 were reported by MiXCR, we would simply join them together as variable region. For sequences whose FR1 to FR4 regions were not completely reported by MiXCR, we extracted them using our algorithm: I) Reads which can not be merged by MiXCR were discarded; II) The beginning of variable region was acquired by pairwise alignment between germline reference of V genes and the column named “targetSequence” reported by MiXCR(The function *pairwise2.align.localms* from Python *Bio module* was used with parameters 2, −3, −5, and −2); III) If the column named “refPoints” in MiXCR recorded the region of FR4, we would use it instead of aligning “targetSequence” to J gene to find the end of FR4.

### Calculation of gene usage diversity

The Shannon index was used to show the diversity of gene usage of public clones. However, as the number of V genes influences the diversity largely, we used the maximum of diversity with particular number of V genes to normalized the diversity. The function to calculate the normalized diversity is shown below. When different donors use totally different V genes, the normalized diversity equals one.

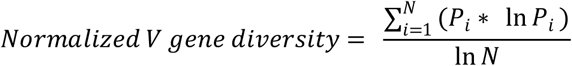

*P*_*i*_ means the frequency of specific V gene, *i* means the order of V gene, and *N* means the total number of V gene.

### Calculation of sequences identity

Both nucleotide and amino acid sequences were aligned by Clustal W 2.1^53^ from the Python module named Bio. Gaps at the beginning and ending of aligned sequences were removed and the percentage of matched bases was defined as identity.

### Multiple sequence alignment for anti PD-1 antibodies

Variable region sequences with the same CDR3aa of Camrelizumab were extracted from each sample and grouped according to the VJ recombination and CDR3nt. Amino acid sequences of groups with most reads in each individual were used for multiple sequence alignment by Clustal W 2.1 with default parameters and visualized by BioEdit.

### Construction of phylogenetic tree

Each phylogenetic tree was generated by the nucleotide sequences of variable regions for antibodies sharing the same CDR3 sequence with MT658807, MT658819, 1-20, MK901823, and KU760937. In addition, the germline V allele of validated neutralizing antibody which was set as the root and validated antibody were also enrolled. Alignments were performed using Clustal W 2.1, and the maximum parsimony trees fitted using DNAMLK by PHYLIP 3.698^54^. Lastly, these phylogenetic trees were displayed and annotated by iTOL^55^.

### Comparison of antibody structure

As some Rep-seq datasets were amplified by Multiplex PCR, variable regions for these sequences were not complete. Thus, sequences lost several bases at the beginning of the FR1 due to the design of primer set were padded by germline sequences from IMGT. Sequences for validated antibodies were downloaded from NCBI. Variable regions without out-frame were used to predict their structures by Repertoire Builder^56^. Then PyMOL was used to calculate RMSD to compare the similarity of antibody structures.

## Acknowledgements

This study was supported by the National Natural Science Foundation of China (NSFC) (31771479) (Z. Z.), NSFC Projects of International Cooperation and Exchanges of NSFC (61661146004), and the Local Innovative and Research Teams Project of Guangdong Pearl River Talents Program (2017BT01S131). We thank Jun Chen from MOE Laboratory of Biosystems Homeostasis & Protection and Innovation Center for Cell Signaling Network, College of Life Sciences, Zhejiang University for the valuable comments, discussions, and suggestions.

## Author Contributions

Y. Z., H. Z., Y. Z., C. L., X. Y., Y. Z., Y. C., Y. Z., J. W., C. W., C. M., and S. C. collected the datasets and performed the bioinformatics analyses. Y. Z. and Q. X. developed and implemented the RAPID platform. M. W., Q. W., H. Tang., W. X., and J. G. collected the samples and conducted the biological experiments. M. W. and S. G. prepared the libraries and ran the Illumina sequencing. C. L. coordinated the project. X. Y. and Z. Z. conceived the project. All authors were involved in the manuscript writing.

## Competing of interests

The authors declared no competing financial interests.

---

**Supp. Fig. 1.**
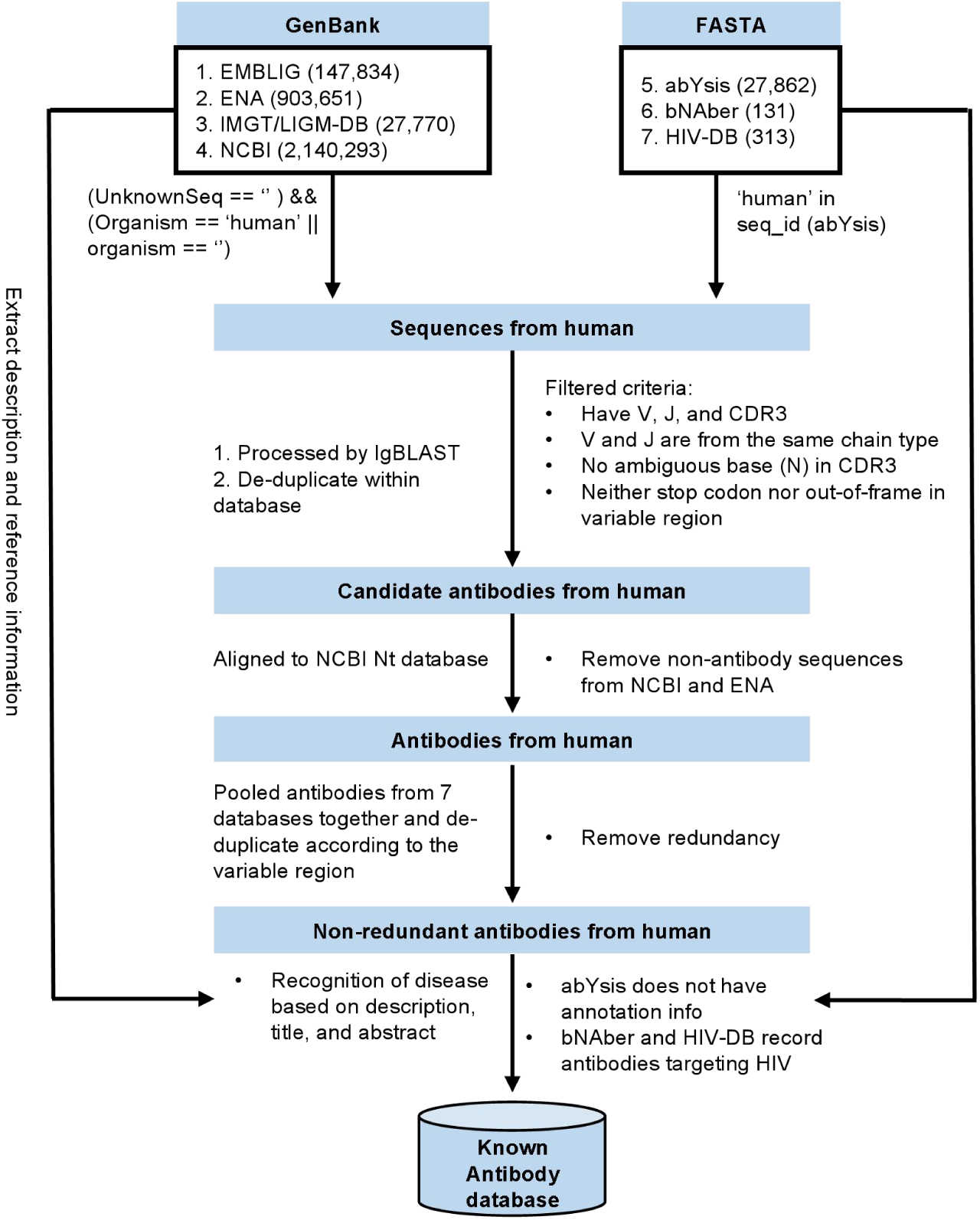
Workflow of known antibody database construction. The first two boxes record the total number of sequences downloaded from 7 databases with Genbank and FASTA formats. Each procession on sequences is marked near arrow between intermediate results.

**Supp. Fig. 2.**
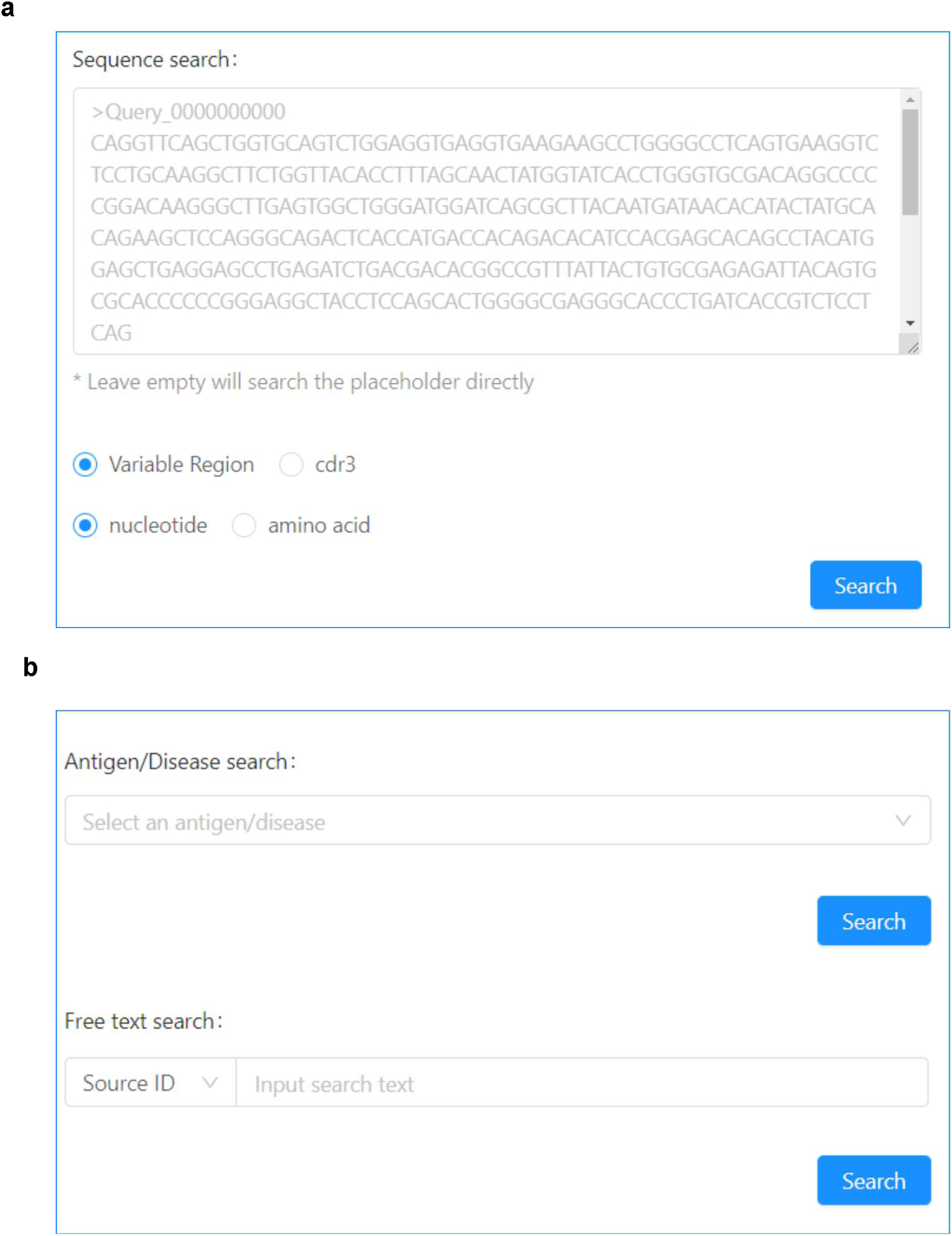
Sequence and text search functions of RAPID. **(a)** The function of nucleotide and amino acid sequences search for both variable region and CDR3. **(b)** Known antibody search based on text such as antigen/disease and source id.

**Supp. Fig. 3.**
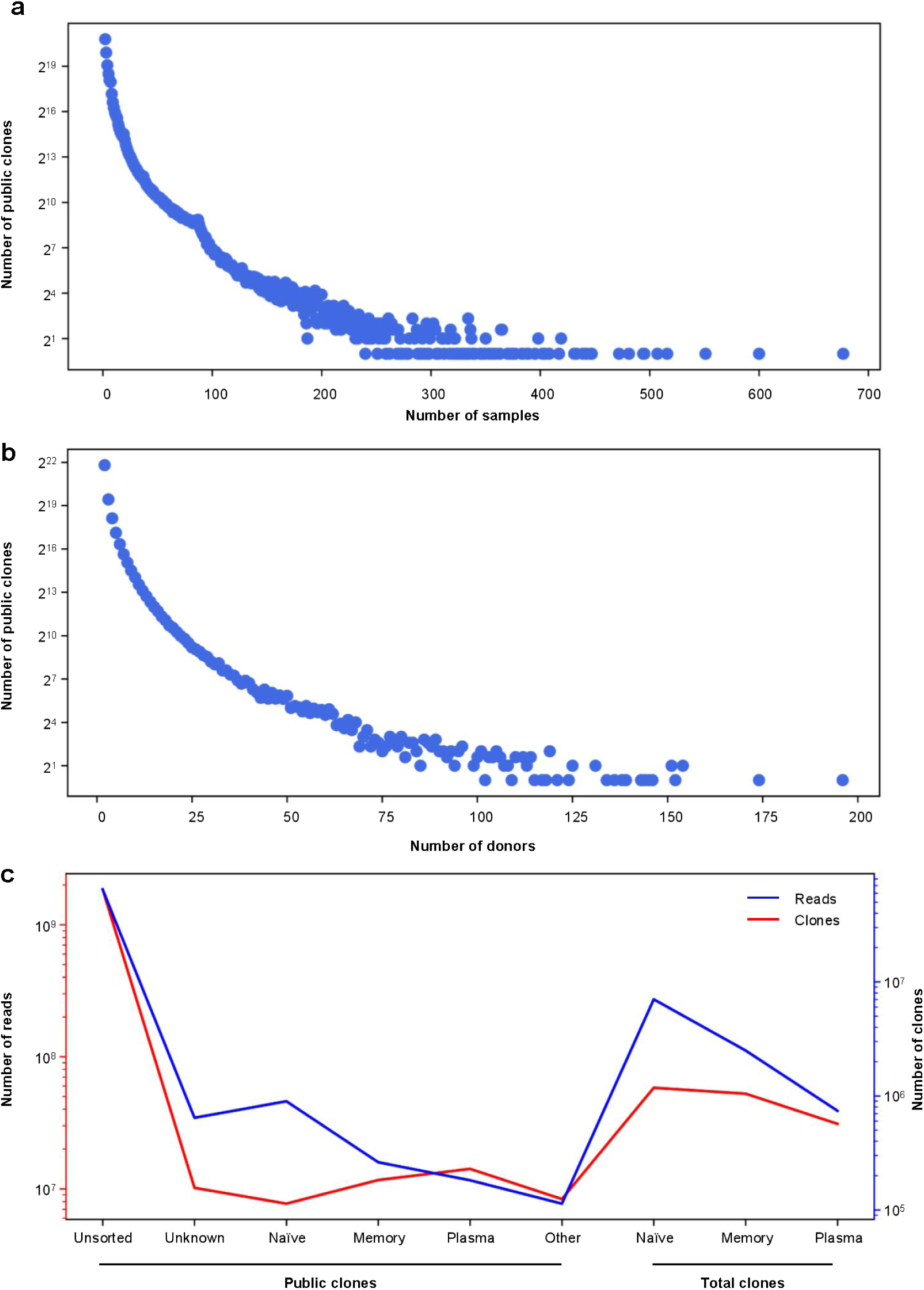
Basic information of public clones. The number of public clone changed with the number of samples **(a)** and donors **(b)**. The Y-axis were logarithmically converted with base 2. **(c)** The number of reads and clones from different sources. Sources of clones were defined based on types of B cell including naïve, memory, plasma, PBMCs, other (particular antigen-specific B cells), and unknown. The Y-axis were logarithmically converted with base 10.

**Supp. Fig. 4.**
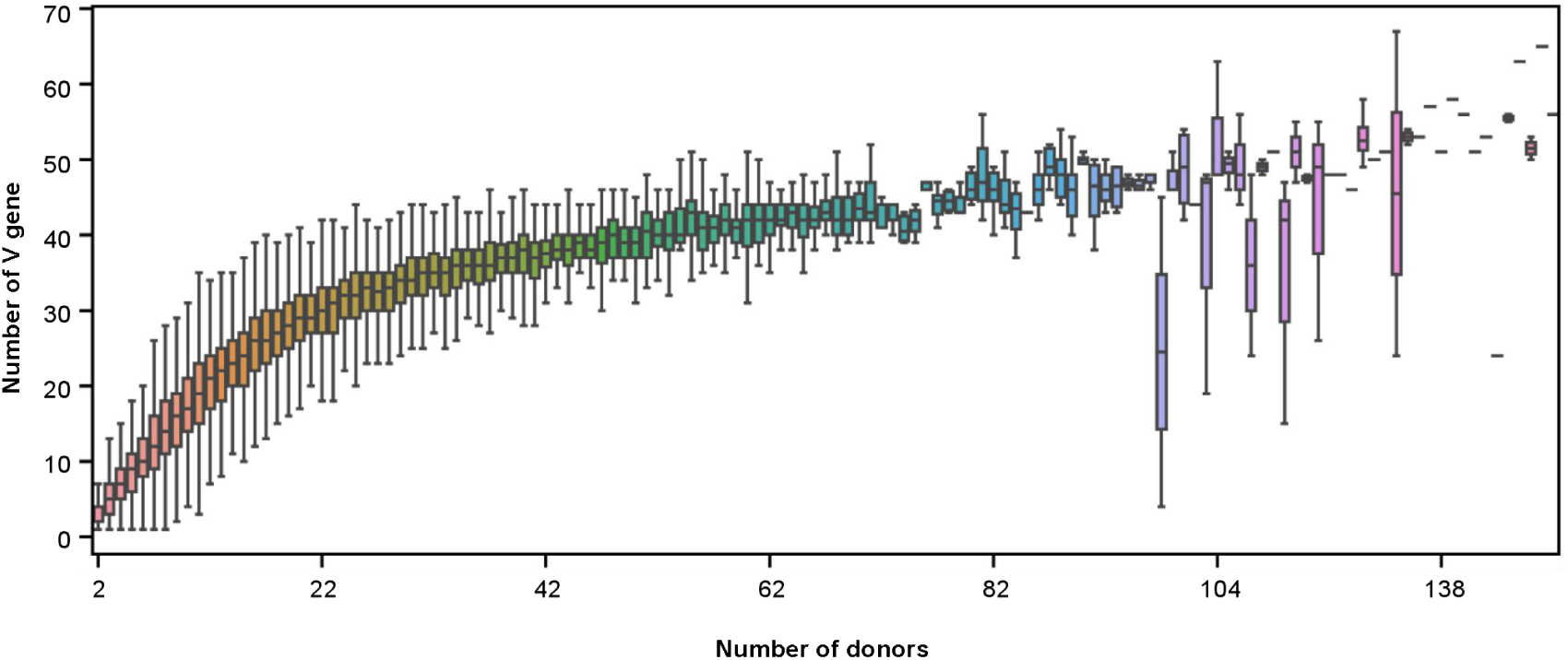
The number of V genes for public clones shared by different number of donors. The number of donors of public clones is discrete.

**Supp. Fig. 5.**
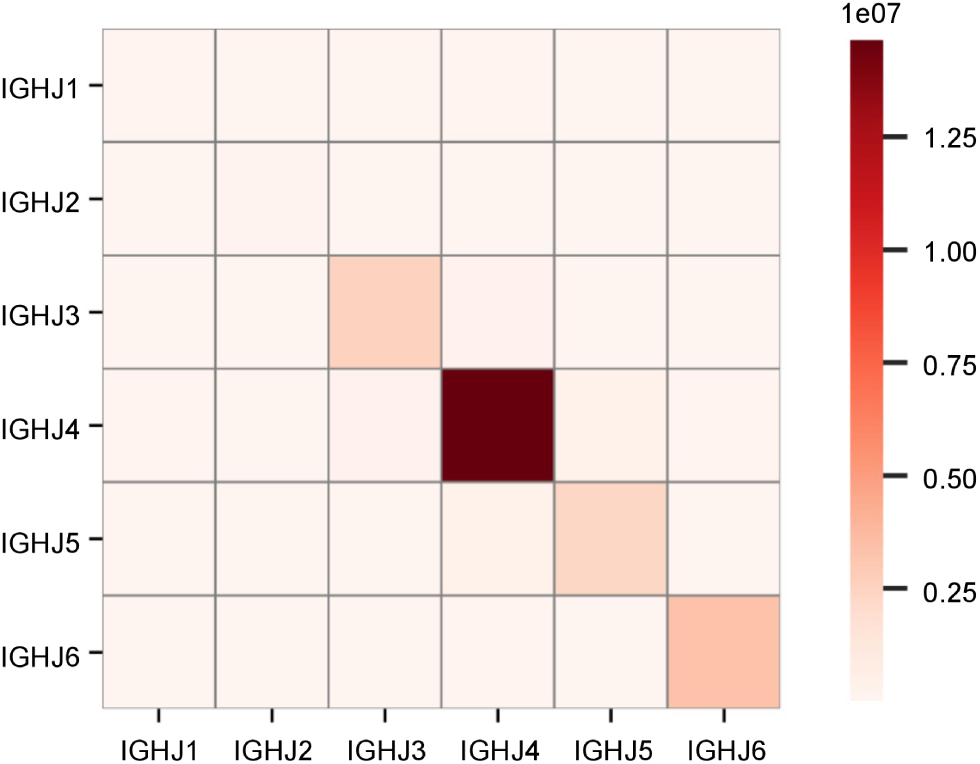
The substitution frequencies of J genes with the same CDR3aa among different donors. The darker the color, the higher the substitution frequency.

**Supp. Fig. 6.**
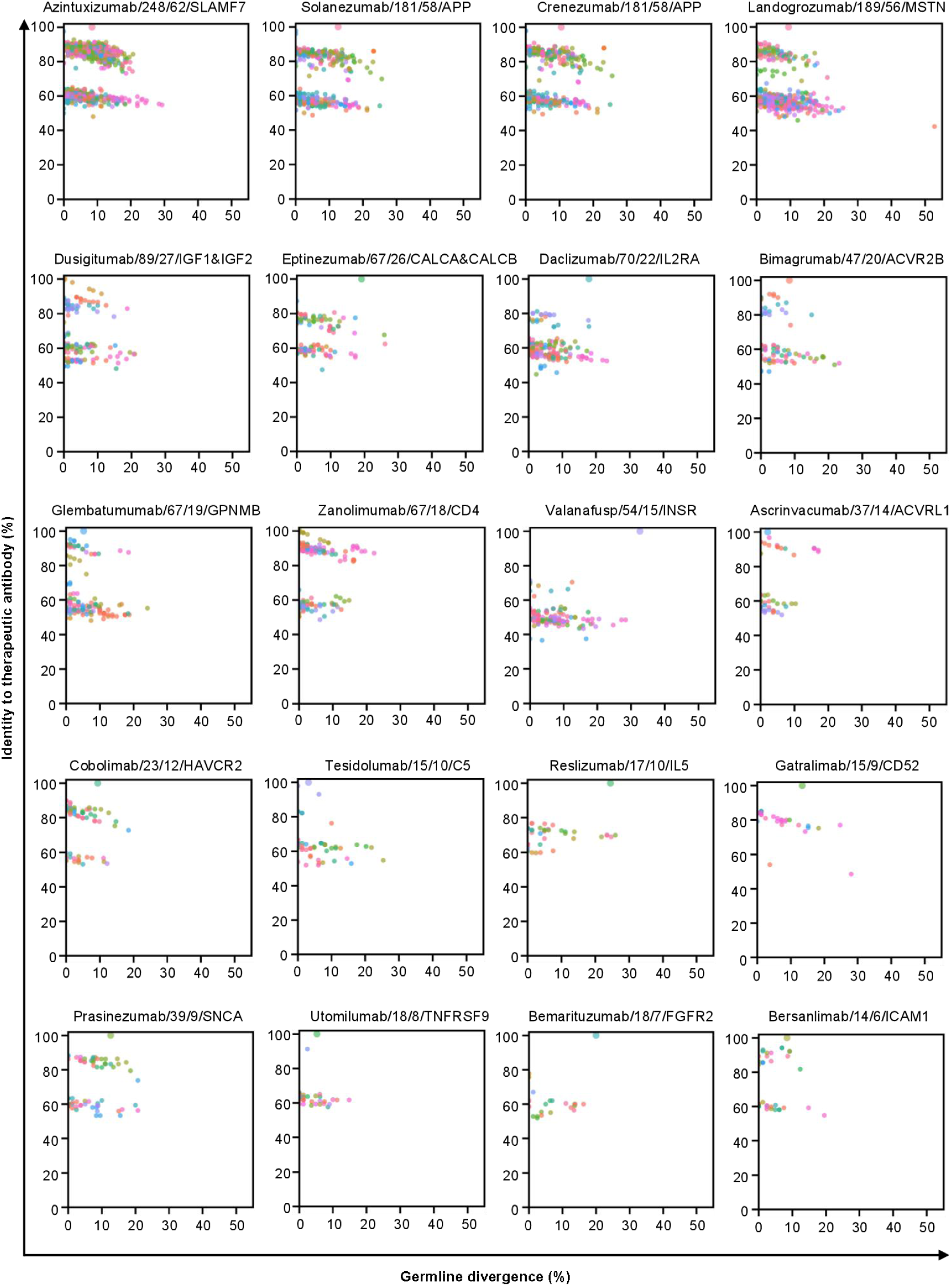
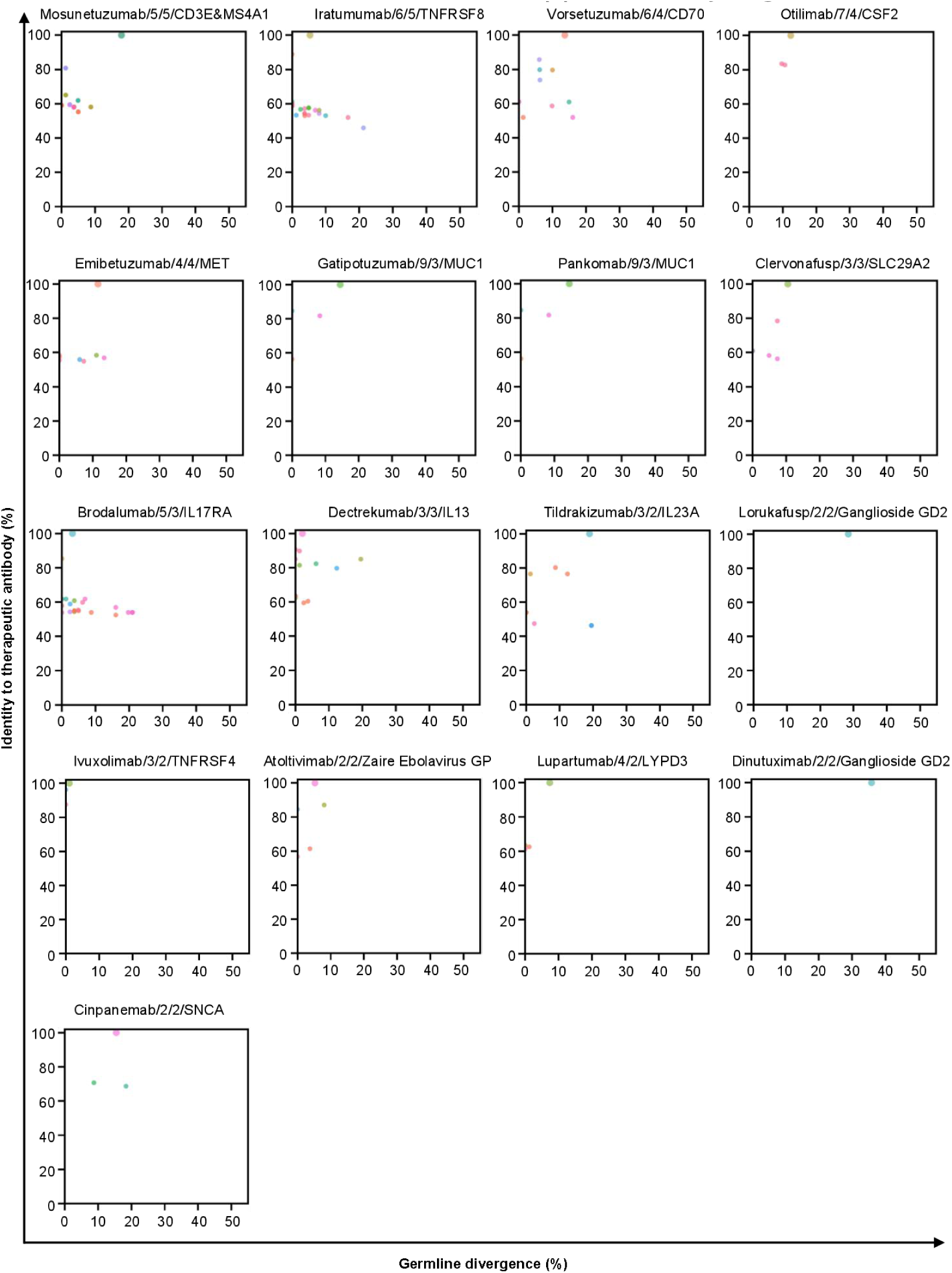
Identity of variable regions from FR1 to FR3 between therapeutic antibody and public clones. The X-axis means the divergence to germline reference and the Y-axis means the sequences identity. Different V genes are filled in different colors. Titles for subfigures separated by forward slash include inn id of therapeutic antibody, the number of samples and donors with such CDR3aa, and target of therapeutic antibody. Dots of therapeutic antibodies are larger than that of clones identified from Rep-seq datasets. Sub-figures are sorted according to the number of donors.

**Supp. Fig. 7.**
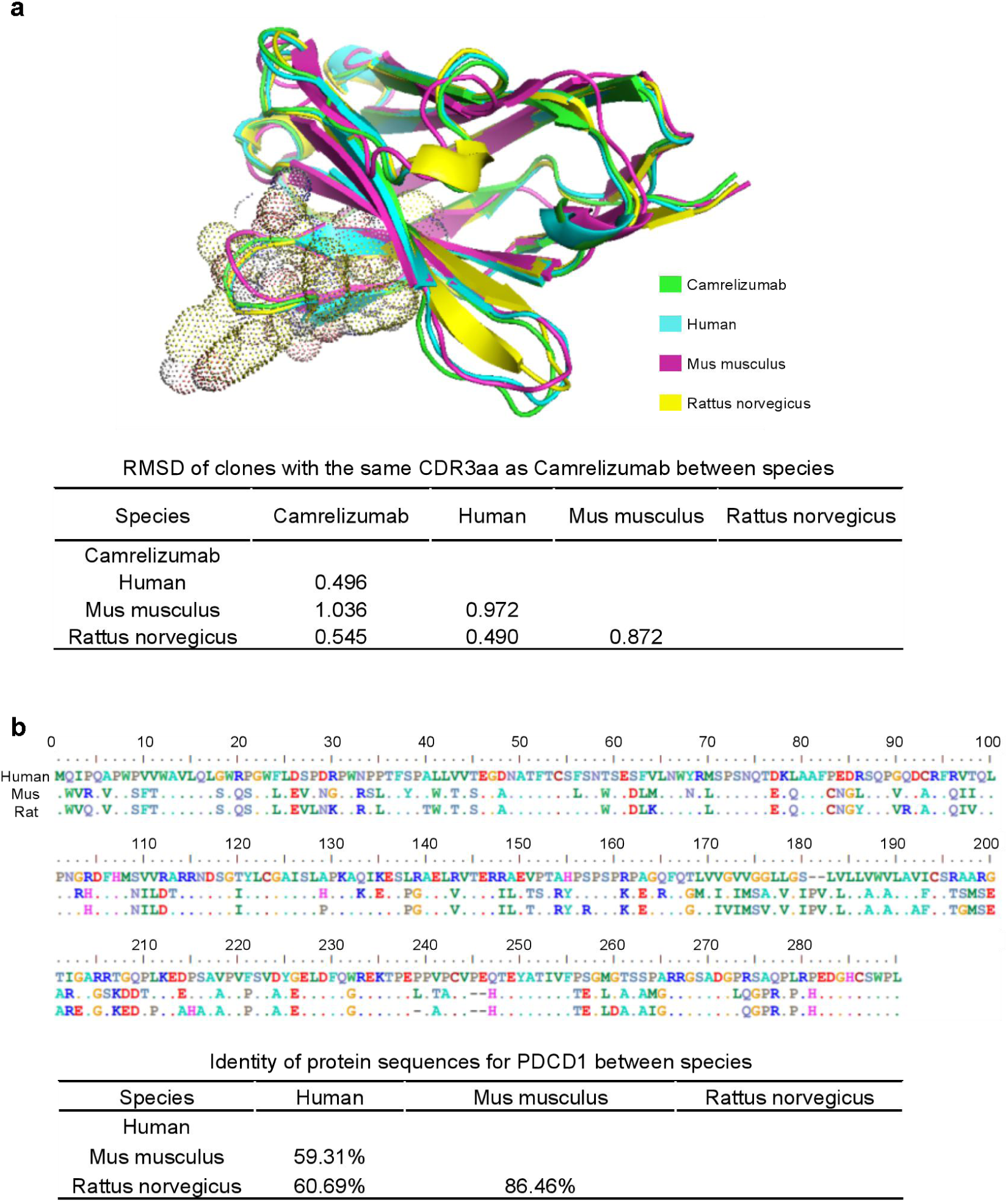
Structure and sequence similarity of anti-PD1 clones and PDCD1 from different species. **(a)** Structure similarity of anti-PD1 clones. The upper panel stands for structures of Camrelizumab and clones from Human, Mus musculus, and Rattus norvegicus. Table in the lower panel records the RMSD of structures between paired species. **(b)** Identity of protein sequences for PDCD1 from human, Mus musculus, and Rattus norvegicus. The upper panel shows the multiple sequences alignment for them and the lower panel shows the sequences’ identity.

**Supp. Fig. 8.**
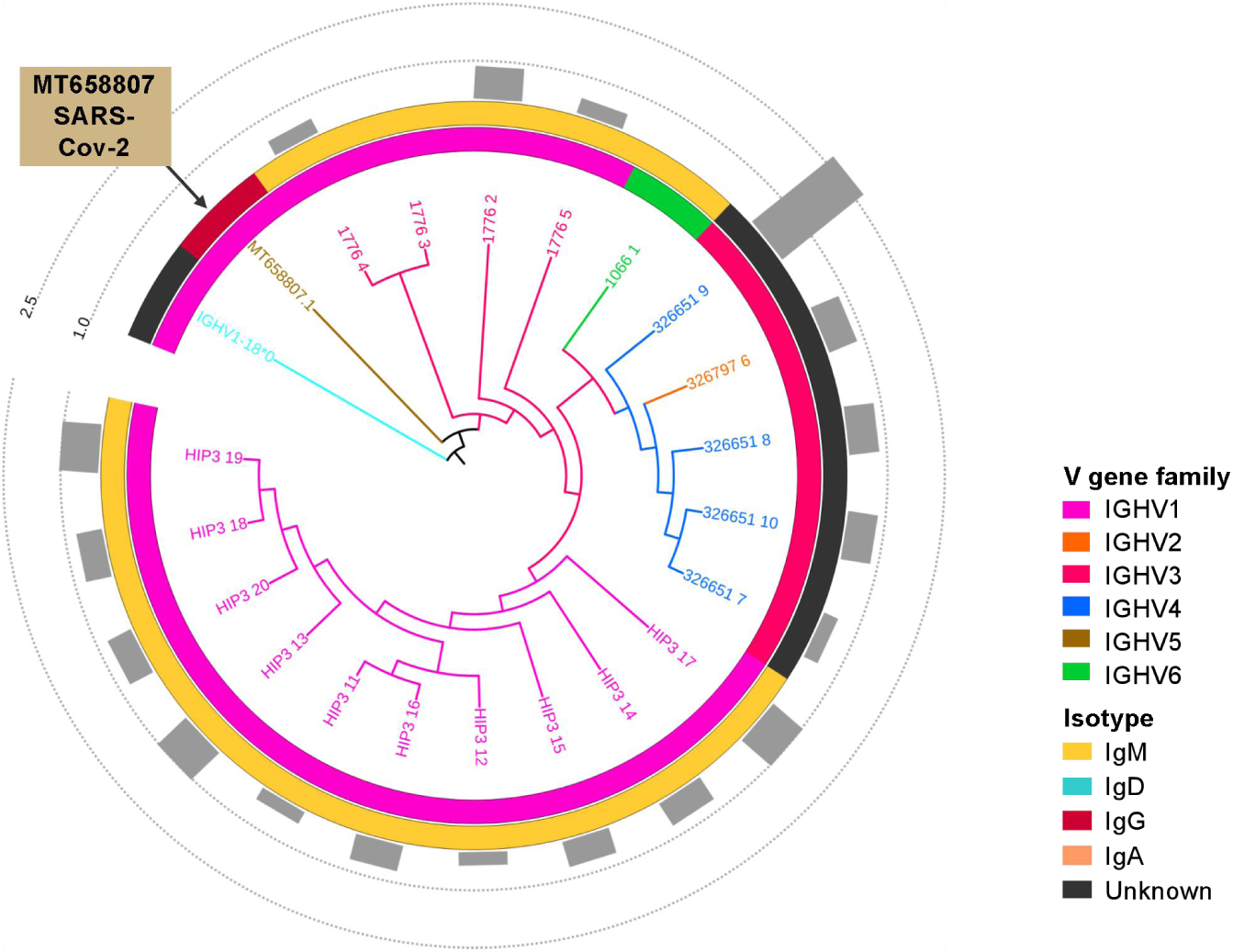
Maturation pathway of clones with the same CDR3aa of MT658807. Variable region sequences with the same CDR3aa as MT658807 were extracted and compared with MT658807 and its’ germline reference. The germline reference was chosen as root of phylogenetic tree and MT658807 is marked by arrow. The cluster map contains four layers including similarity of sequences (the sequences extracted from the same donor were marked with the same color), V gene family, isotype, and somatic hypermutation rate from inner to outer.

**Supp. Fig. 9.**
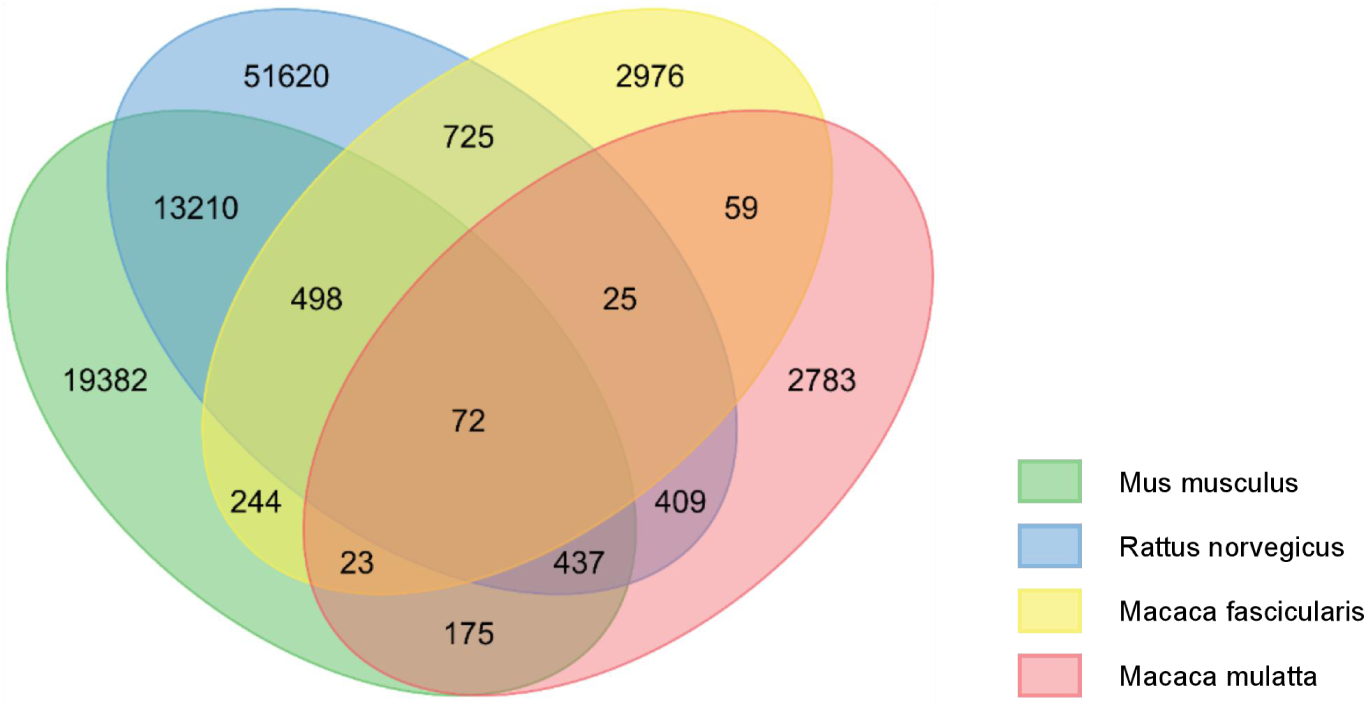
Overlap of public clones shared by other species.

## Notes

### Competing Interest Statement

The authors have declared no competing interest.

### Summary of Updates

Supplementary figure 7 revised; Supplemental files updated.

